# Myxococcus xanthus fruiting body morphology is important for spore recovery after exposure to environmental stress

**DOI:** 10.1101/2023.05.19.541530

**Authors:** Dave Lall, Maike M. Glaser, Penelope I. Higgs

**Author notes:** Address correspondence to Penelope I. Higgs,.

## Abstract

Environmental microorganisms have evolved a variety of strategies to survive fluctuations in environmental conditions, including production of biofilms and differentiation into spores. *Myxococcus xanthus* are ubiquitous soil bacteria that produce starvation-induced multicellular fruiting bodies filled with environmentally resistant spores (a specialized biofilm). Fruiting bodies are thought to facilitate the *M. xanthus* social life cycle by ensuring spores can germinate *en masse* into a productive feeding community. Isolated spores have been shown to be more resistant than vegetative cells to heat, ultraviolet radiation, and desiccation, but it is unknown whether assembly of spores into a fruiting body provides additional protection from environmental insults. We developed a high-throughput method to compare the recovery (outgrowth) of distinct cell types (vegetative cells, free spores, and intact fruiting bodies) after exposure to ultraviolet radiation or desiccation. Our data indicate haystack-shaped fruiting bodies protect spores from extended UV radiation but do not provide additional protection from desiccation. Perturbation of fruiting body morphology strongly impedes recovery from both UV exposure and desiccation. These results hint that the distinctive fruiting bodies produced by different myxobacterial species may have evolved to optimize their persistence in distinct ecological niches.

**IMPORTANCE:** The myxobacteria are environmentally ubiquitous social bacteria that influence the local microbial community composition. Understanding how these bacteria are affected by environmental insults is important in predicting how microbial biogeochemical cycling is affected by climate change. When starved, myxobacteria produce multicellular fruiting bodies filled with spores. As spores are resistant to a variety of environmental insults, it has long been held that the fruiting body evolved to ensure group germination into a productive feeding community. Using the model myxobacterium, *Myxococcus xanthus*, we demonstrate that the haystack-shaped fruiting body morphology enables significantly more resistance to UV exposure than the free spores. In contrast, fruiting bodies are slightly detrimental to recovery from extended desiccation, an effect that is strongly exaggerated if fruiting body morphology is perturbed. These results suggest the variety of fruiting body morphologies observed in the myxobacteria may dictate their relative resistance to changing climate conditions.

## INTRODUCTION

Environmental communities of microorganisms play important roles in the global ecosystem by contributing to biogeochemical cycling of carbon, nitrogen, and other essential elements (1, 2). The myxobacteria are a group of highly social predatory or cellulolytic bacteria that are thought to be important for influencing environmental microbial community structure (3–7). Analysis of data from the Earth Microbiome Project determined that the myxobacteria are among the most widely distributed prokaryotic orders on Earth (8). They have been identified in a wide variety of terrestrial and marine environments (9), including some extreme environments, such as the Sahara with a high UV index and long periods of desiccation (10). Thus, understanding how myxobacteria are able to persist in environments is important to fully understand how microbial communities will be affected by, and consequently contribute to, climate change (11, 12).

Most Myxobacteria have the ability to produce dormant spores that display increased resistance to environmental insults such as UV, desiccation, and heat (13). Interestingly, myxobacterial spores are typically produced inside of macroscopic fruiting bodies produced at the culminating stage of an elaborate multicellular developmental program (13). Different myxobacterial genera produce distinctive fruiting bodies, with heights in the range of 0.1-1 mm and morphologies ranging from simple mounds containing spores embedded in extracellular matrix (ECM) to elaborate tree-like structures in which spores are housed in rigid, thick-walled sporangioles (13, 14). It has long been held that it is the spore state which enables survival under unfavorable conditions, while the function of fruiting bodies is merely to keep the spores together to allow dispersal and/or germination *en masse* upon return to favorable conditions (13). The latter hypothesis is suggested because most of the myxobacteria obtain nutrients in a process that is facilitated by collective secretion of degradative enzymes to break down external macromolecules.

The best characterized of the myxobacteria is *Myxococcus xanthus*. In the vegetative state, cells obtain nutrients by cooperative predation mediated by collective secretion of antibiotics, proteases, lipases and other lytic enzymes (15, 16). Thus, growth rate is proportional to population density (17). Upon nutrient limitation*, M. xanthus* communities form simple haystack-shaped, soft, fruiting bodies each consisting of approximately 10^5^ metabolically quiescent spores embedded in a matrix of exopolysaccharides, proteins, lipids and eDNA (18–21). Relative to the vegetative cells, *M. xanthus* spores have been shown to have enhanced resistance to UV, desiccation, heat up to 60 °C for 60 min, enzymatic digestion, detergents and sonic disruption (22–24). These resistance properties fall far short of endospores produced by common environmental Firmicutes (25).

Here, we explore the hypothesis that myxobacterial fruiting bodies provide additional protection to the spores from environmental insults. We first developed a high-throughput method to assay community recovery after exposure to two common environmental insults, UV exposure and desiccation. We use this method to confirm that free spores are more resistant than vegetative mats to UV exposure, but the fruiting body community can recover after ∼8-fold longer exposure. An *M. xanthus* mutant strain that produces only shallow fruiting bodies shows no advantage over free spores. In contrast, relative to the free spores, wild-type fruiting bodies provide little additional protection to short term (<1 week) desiccation and hinder recovery from extended (> 2 week) desiccation. However, production of shallow fruiting bodies significantly impedes recovery compared to free spores, likely due to severe dehydration of the fruiting body matrix material. These studies have important implications for environmental community composition in the face of more frequent and extreme environmental fluctuations arising from climate change.

## MATERIALS AND METHODS

### Strains and media

The following *M. xanthus* strains were used in this study: DZ2 (26), PH1054 [DZ2 1′*espA* 1′*espC* 1¢(*redCDEF) todK::miniTn5 8846*; here termed Shallow Fruiting (SF) mutant], PH2036 [DZ2 *attB*::Pr_pilA_-*mCherry* (pFM16 (27); Kn^R^)], and PH2037 (PH1054 *attB*::pFM16). Strains were grown under vegetative conditions in CYE broth [1% Casitone, 0.5% yeast extract, 10 mM 3-(*N-*morpholino) propanesulfonic acid (MOPS) pH 7.6, 4 mM MgSO_4_) (26) on an orbital shaker at 220 rpm and 32 °C. Strains were recovered from permanent stocks on CYE agar plates (CYE, 1.5% agar), containing kanamycin at 100 ug ml^-1^ as necessary, and grown at 32 °C.

### Bacterial growth curve measurements

To generate flask culture growth curves, cells were inoculated into 15 mL of CYE broth and grown overnight at 32 °C with shaking at 220 rpm. Cells were subcultured 1:100 into 45 mL fresh CYE broth in 500 mL Erlenmeyer flasks and incubated as above. Every four hours, 1 mL of culture was withdrawn and absorbance at 550 nm (A_550_) was recorded in a spectrophotometer (Ultrospec 2100, Amersham Biosciences) using cuvettes with 1 cm pathlength (Table 1: flask growth, spectrophotometer measurement), or by placing 0.2 mL in triplicate wells of a 48-well plate followed by absorption measurement in a Tecan Spark 10M plate reader (Table 1: flask growth, plate reader measurement). Once samples reached A_550_ > 1, they were diluted in CYE before absorbance readings.

For growth curve analysis of broth cultures in 48-well plates (CytoOne, USA Scientific), overnight cultures were diluted to an A_550_ of 0.035 in fresh CYE and 200 μL was seeded into triplicate wells. Plates were incubated in a Tecan Spark 10M for 72 hours with continuous orbital shaking (216 rpm) at 32 °C, and the A_550_ was recorded automatically every 2 hours (Table 1: plate reader growth and measurement). Each growth curve was blanked to the respective initial (T=0) A_550_ reading. Final growth curves were plotted as the average and associated standard deviation of A_550_ values recorded from three independent biological replicates each containing three technical replicates, versus time.

The length of the lag phase was measured as the length of time to reach A_550_ of 0.02 which was determined by solving for time using the slope equation from a trendline generated using three data points surrounding the 0.02 A_550_ value. Doubling times (t_d_) were calculated using the equation t_d_ = ln 2 / μ, where μ is the growth rate. Growth rate was calculated as μ = (ln A_2_ – ln A_1_) / (t_2_ – t_1_), where t_1_, A_1_ and t_2_, A_2_ correspond to values in the exponential growth phase (28). Onset of stationary phase was taken as the time corresponding to the peak A_550_ value recorded.

### Generation of cell states

To generate vegetative mats, overnight cultures were diluted to 0.035 A_550_ in fresh CYE and 200 μL was seeded in triplicate wells of a 48-well plate (CytoOne, USA Scientific), followed by incubation for 24 hours at 32 °C without shaking. Fruiting bodies were generated in submerged culture (29). Briefly, the CYE over-layer was removed from vegetative mats, replaced with 200 μL of sterile MMC starvation media (10 mM MOPS, pH 7.6, 2 mM CaCl_2_, 4 mM MgSO_4_), and further incubated at 32 °C without shaking for 120 hours. To generate free spores, submerged cultures were generated in 100 mm petri dishes as above, except 16 mL volumes were used. Five days post starvation, fruiting bodies were harvested, pelleted (4 600 x g, 10 min, RT), resuspended in 0.5 ml sterile water, and spores were dispersed by mild sonication at output 10%, 0.5 sec on/0.5 sec off for a total of 23 seconds (Branson Sonifer 250). Spores were diluted ∼30-fold in sterile water and 200 µL volumes were added to triplicate wells of a 48 well plate. Spores were allowed to settle onto the surface of the well for 60 min at RT. Sporulation assays were performed as described previously (29).

For outgrowth analysis of the vegetative mats, fruiting bodies, or free spores, samples were generated in triplicate wells in a 48 well plate, the respective overlay media was replaced with 200 μL fresh CYE, and plates were incubated in the plate reader with continuous orbital shaking (216 rpm) at 32 ^°^C for 72 hours. A_550_ values were automatically recorded every 2 hours. CYE media served as the blank. Growth curves were plotted as described above. Final outgrowth curves were the average and associated standard deviation of nine growth curves generated from three independent biological replicates each containing triplicate technical replicates.

### UV stress assay

Vegetative mats, fruiting bodies, or free spores were generated in 48 well plates as described above, with one plate used per cell state. Wild type and SF mutant strains were added to 21 wells each; 3 wells served as cell-free blanks. Overlay media from all wells was replaced with 50 μL of sterile water, and triplicate wells for each strain were exposed to 0, 1, 2, 4, 8, 16, or 32 minutes of 306 nm UV supplied by an inverted transilluminator rated at 8000 μw/cm^2^ (Model FB-TI-88A, Fisher Scientific, USA) that was placed 10 cm above the uncovered plate. Aluminum foil was used to shield the remaining wells. Following UV exposure, an additional 50 μL sterile water and 100 μL of fresh 2X CYE were added to each well, and outgrowth was measured in a plate reader as described above.

### Desiccation Assay

For desiccation of vegetative mats, wild-type and SF mutant strains were each seeded in 3 wells (48 well plate), and the plate was incubated at 32 °C without shaking. This process was repeated 16, 24, 28, 30, 31, and 32 days later. Twenty-four hours after the respective seeding times, CYE media was removed and plates were incubated at 32 °C. Thus, triplicate wells of each strain experienced 32, 16, 8, 4, 2, 1, or 0 days of desiccation. 33 days after the initial seeding, 200 μL of fresh CYE was added to all wells to induce outgrowth. 3 wells served as cell-free blanks. Outgrowth was measured in a plate reader as described above. Desiccation of fruiting bodies and free spores followed the same pattern, except cells were seeded 6 days in advance of the respective desiccation start times. Free spores were generated from 100 mm petri dishes as described above.

### Determination of time to recover after insult

Recovery curves were plotted as described for growth curves above. Final recovery curves were plotted as the average A_550_ and associated standard deviation from three independent biological replicates each containing three technical replicates, versus time. An A_550_ of 0.2 was selected as a reference indicating early exponential growth. Recovery time was calculated using the slope equation from a trendline derived from four data points surrounding A_550_ = 0.2. The relative fold recovery for each cell state was expressed as treated recovery time (t_T_) / untreated recovery recovery (t_UT_). All graphs and statistical analyses were generated in GraphPad Prism (v. 9.5.1). Statistical significance of relative recovery times was determined by two-way ANOVA followed by Tukey’s multiple comparison test (\*\*\*\**p* <0.0001). Figures were generated in Affinity Designer 1.10.6.

### Imaging of fruiting bodies, vegetative mats, and free spores

To image vegetative mats, fruiting bodies, and desiccated fruiting bodies, the respective cells states were generated μ-Dishes ^35^ ^mm, high^ (ibidi Technologies; hereafter ibidi dishes), except 2.1 mL volumes of seed cultures or starvation buffer was used. Images were captured using a Leica M80 stereomicroscope and a Leica DMC2900 camera (50X magnification), or a Zeiss Axio Imager. M1 microscope and Cascade 1K camera (100-1000X magnifications). Free spores were generated as described above, and 10 μL aliquots were spotted onto agar pads (30).

For confocal imaging of wild-type and SF mutant fruiting bodies, strains PH2036 and PH2037 (expressing mCherry from the constitutive pilA promoter in DZ2 and PH1054 backgrounds, respectively) were developed in ibidi dishes for 48 hours.. Fruiting bodies were imaged using a Leica TCS SP8 inverted confocal microscope. mCherry fluorescence was detected using a 552 nm wavelength laser (5% power) for excitation, a 585-630 nm emission spectra, and gain of 750 V. Fruiting bodies were imaged from the base to the top with a step size of 1 μm, and line average of 4 at 1024 x 1024 resolution.

To determine fruiting bodies heights, wild type (DZ2) and SF mutant (PH1054) strains were induced to develop in ibidi dishes for 5 days. Fruiting bodies were stained in MMC supplemented with 2 mM MnCl_2_ and 15 μl of Fluorescein conjugated Concanavalin A (5mg/ml; solubilized in 0.1 M NaHCO_3_ pH 8.3, Karlsruhe, Germany), and incubated for 30 min at RT in the dark. The stain was removed and fruiting bodies were washed twice with 1 ml MMC, 2 mM MnCl_2_. For microscopy, 2 ml of MMC were added back to the cells. The fluorescein signal was detected using a 552 nm wavelength laser (15% power) for excitation, a 494-572 nm emission spectra with an inverted TCS-SP5 confocal microscope (Leica, Bensheim, Germany). Fruiting bodies were imaged from the base to the top with a step size of ∼0.9 μm. Image data and fruiting body heights were processed by using IMARIS software package (Bitplane AG, Zurich, Switzerland) or Volocity Software (Perkin Elmer).

## RESULTS

### Growth characteristics of *M. xanthus* cultivated in 48-well plates

We first set out to generate a high-throughput method to monitor growth of *M. xanthus* wild-type strain DZ2 using a plate reader to measure growth in 48 well plates. To understand how variables such as aeration, inability to dilute samples of dense cultures, and absorbance pathlength affected the apparent growth characteristics, we compared data obtained from growth of the wild type strain under standard broth culture conditions to that obtained by the plate reader. *M. xanthus* growth and measurement in standard conditions produced a reproducible growth curve with distinct lag, log, stationary, and death phases (Fig. S1). The wild type doubling time was calculated as 5.3 hours and peak cell density was measured at 6.0 A_550_ (Table 1).

**Table 1.**
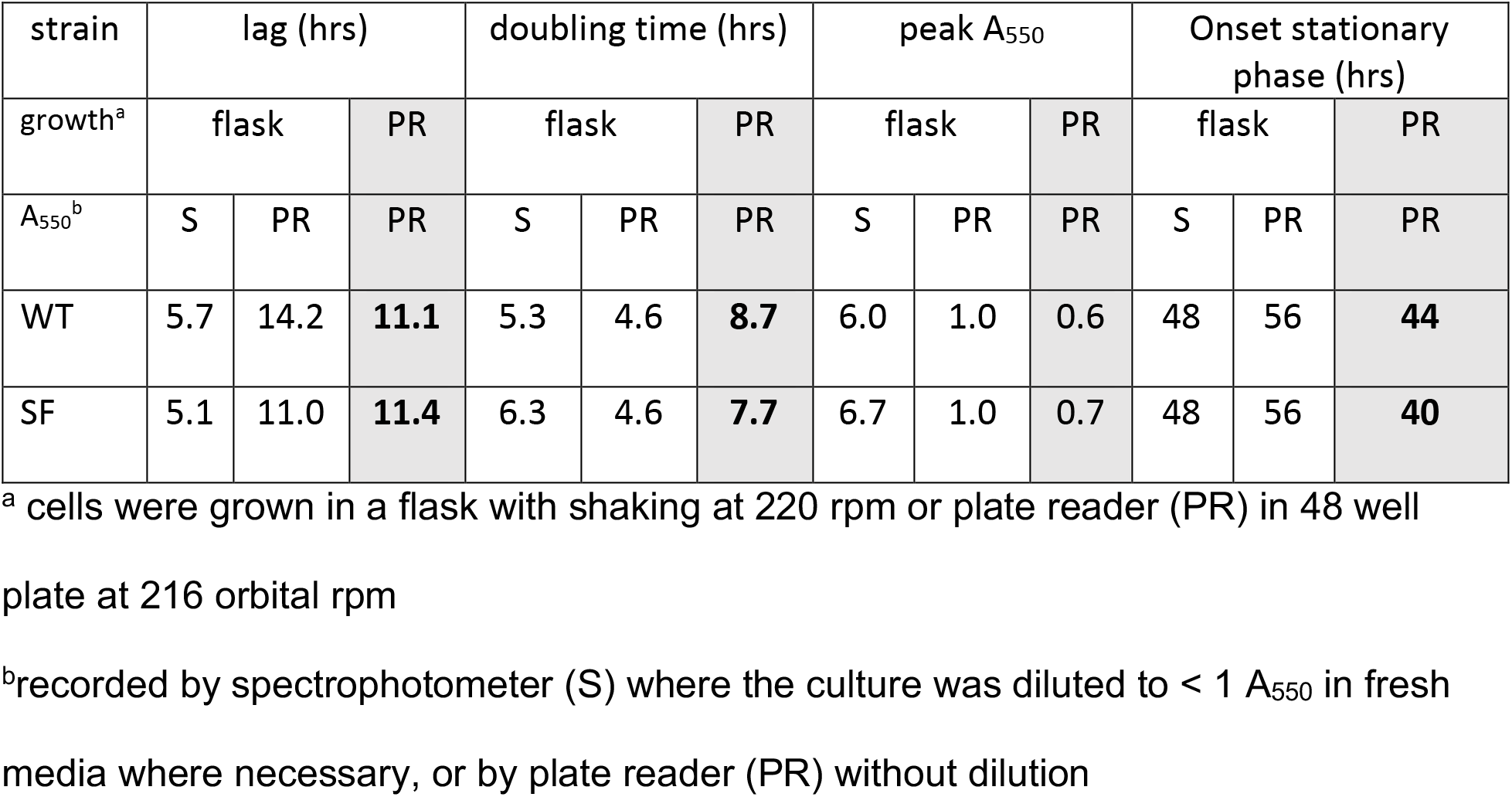
Growth characteristics of *M. xanthus* strains grown in 48-well tissue culture plates.

When the absorbance of the flask cultures was instead measured undiluted in a plate reader, the overall apparent A_550_ was reduced (Fig. S1B), likely because the pathlength of the absorbance measurement is 0.26 cm compared to 1 cm in the spectrophotometer. Analysis of these data lead to aberrations in the calculated doubling time, lag time, and onset of stationary phase (Table 1). This may be because at higher cell densities, the correlation between absorbance and cell number is outside of the dynamic range.

With these parameters established, we next ascertained how culturing *M. xanthus* in 48-well plates affected growth characteristics. We observed that growth of the wild type in wells shortened the lag phase by approximately 3 hours and increased the doubling time approximately 4 hours (Fig. S1B and Table 1). Furthermore, onset of the saturation phase occurred 12 hours earlier and a death phase could be observed within the 72 hours for which data was collected (Fig. S1B and Table 1). These results suggest that cells cultured in wells may be limited by oxygen and/or nutrient availability compared to those growing in flasks.

### Relative outgrowth of *M. xanthus* vegetative mats, free spores, and fruiting bodies

We next took advantage of the plate reader to determine the relative outgrowth of the distinct cell states: vegetative mats, fruiting bodies, or free spores dispersed from fruiting bodies (Fig. 1). To monitor outgrowth of each state, the respective overlay was replaced by fresh nutrient-rich broth, and the plates were incubated in the plate reader using the incubation parameters defined above. We observed that vegetative mats experienced a lag phase of 5.5 ± 0.8 hours, compared to 14 ± 2 and 24 ± 1 for mature fruiting bodies and free spores, respectively (Fig. 2A). The longer lag phase observed by fruiting bodies and free spores reflects the requirement for these cell states to first germinate. Additionally, however, each cell state contains different numbers of initial cells/spores, because approximately 80% of cells lyse during development and not all spores are viable (31). We estimated vegetative mats contain ∼5 x 10^7^ cells per well (31), whereas fruiting bodies contained 6.5 x 10^6^ spores and were estimated to contain ∼2 x10^6^ peripheral rods per well (31)]. We determined free spores were present at approximately 4 x 10^6^ spores per well.

**FIG 1.**
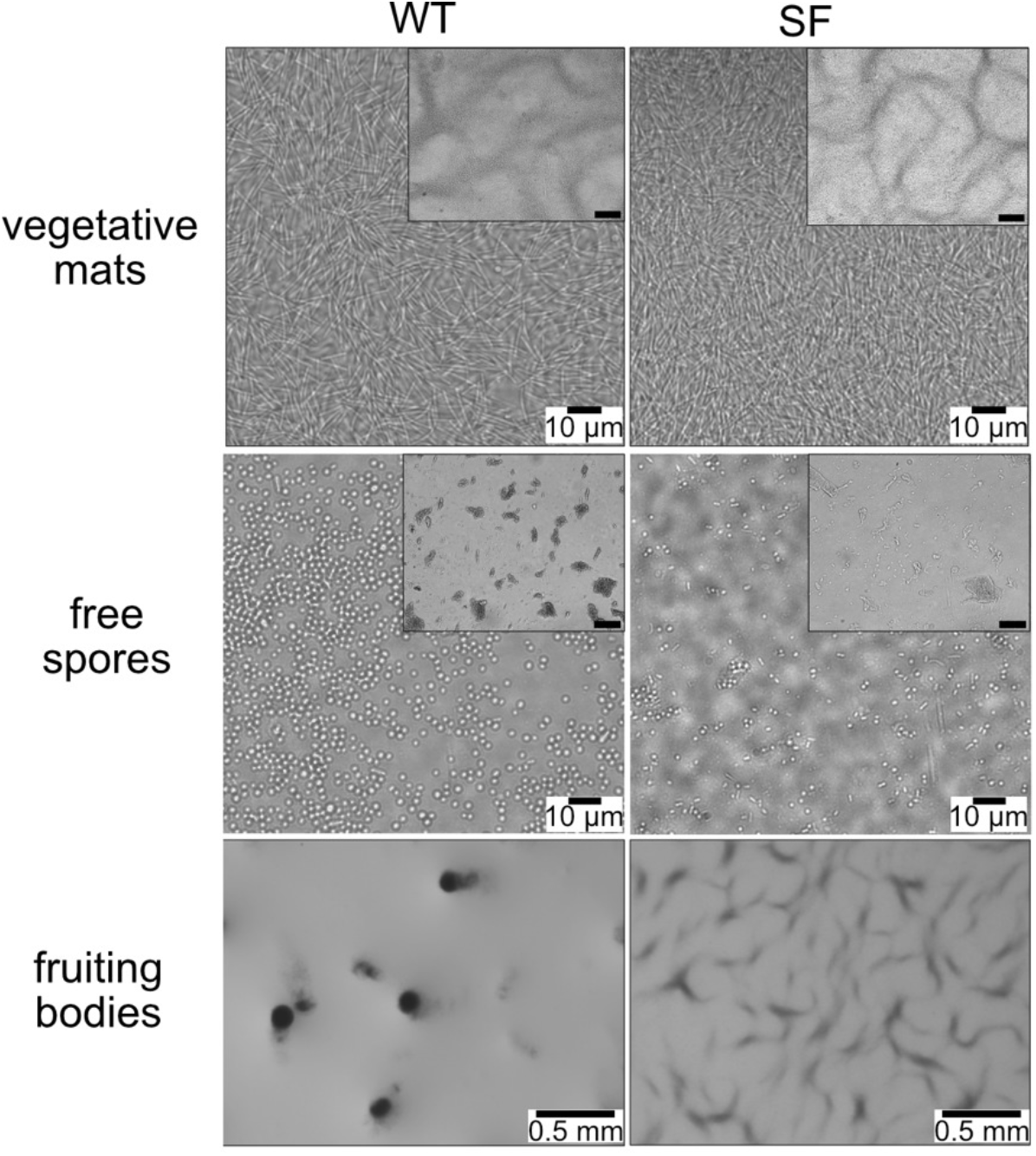
Cell states produced by wild type (WT; strain DZ2) and a shallow fruiting (SF; strain PH1054) strains. Vegetative mats were grown for 24 hours at 32°C, free spores were released from fruiting bodies with mild sonication, and fruiting bodies were induced to develop under submerged culture for 120 hours. Scale bars represent 100 μm unless otherwise indicated.

**FIG 2.**
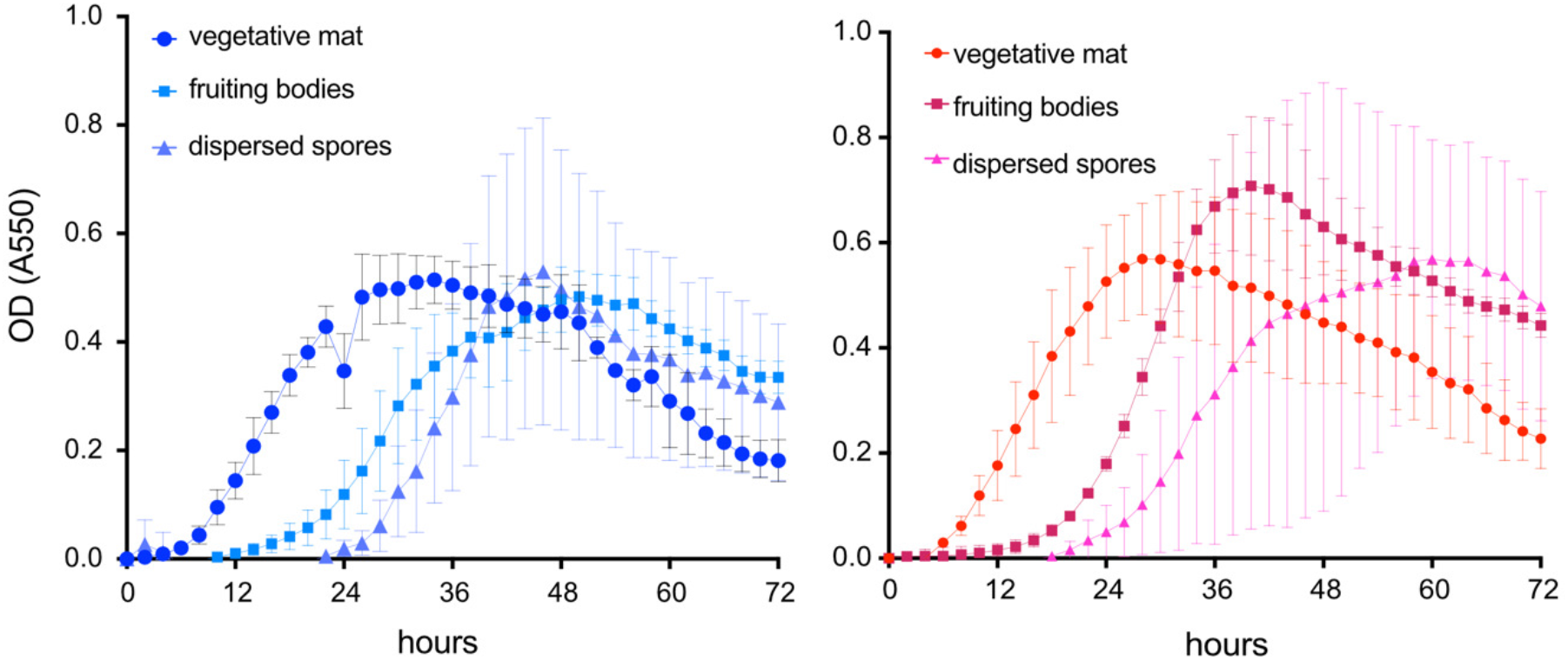
Outgrowth of cell states in the wild-type (DZ2) (A) and shallow-fruiting mutant (PH1054) (B) strains. Vegetative mats (circles), fruiting bodies (squares), and dispersed spores (triangles) were generated in 48 well plates. To induce outgrowth, rich media was added (0 hours), cells were incubated at 32 ^°^C with shaking in a plate reader, and A_550_ was recorded every 2 hours for 72 hours. Data points are the average and associated standard deviation of three independent biological replicates each containing three technical replicates.

### Assembly of spores into fruiting bodies provided additional protection against UV stress

Having established outgrowth characteristics of each cell state, we next examined the relative outgrowth rates after exposure to UV. For these experiments, triplicate wells of vegetative mats, free spores, or intact fruiting bodies were exposed to 0, 1, 2, 4, 8, 16, and 32 min of UV at 306 nm. Outgrowth was then induced and monitored as described above. Analysis of three independent biological replicates indicated the time to recover to early log phase corresponded to increasing length of UV exposure (Fig. S2). These results suggested that the relative recovery period reproducibly corresponded to the fraction of the community surviving UV exposure, rather than to spontaneous mutations leading to variable growth rates. Furthermore, the relative recovery pattern was also observed if cells were instead harvested, serially diluted, and directly spotted on rich media agar plates (Fig. S4). Together, these data suggest relative recovery time was useful as a way to generate quantitative data on the relative tolerance to insult exposure.

To compare the relative ability of the distinct cell states to withstand UV exposure, we ascertained the time for each exposed culture to reach early log phase (apparent A_550_ =0.2). These recovery times were then normalized to the time required for the untreated sample (0 min UV exposure) to reach 0.2 A_550_. The resulting relative recovery times were plotted against minutes of UV exposure for each cell state (Fig. 3). For the wild-type vegetative mats, 1 min of UV exposure resulted in an average 3-fold increase in time for the community to reach 0.2 A550 (Fig. 3). No outgrowth was detected (ND) after exposure to 2, 4, or 8 min of UV. Analysis of the free spore community suggested that after 1 min UV exposure, spores required only a 1.5-fold increased recovery time. Thus, spores provide some protection from UV irradiation, as has been previously demonstrated (22, 24). However, we did not detect recovery of free spores after 2 min or more of UV exposure (Fig. 3). In contrast, recovery of the fruiting body cell state was readily observed, requiring only 2-fold longer recovery time than the untreated sample even after 8 min of UV exposure. No recovery was detected for any cell state after 16 and 32 min of UV exposure (data not shown). These results indicated that while spores do provide protection over vegetative cell mats to shorter intervals of UV exposure, the fruiting body state provides much more protection to extended UV exposure, likely because UV may not effectively penetrate the interior of the fruiting body.

**FIG 3.**
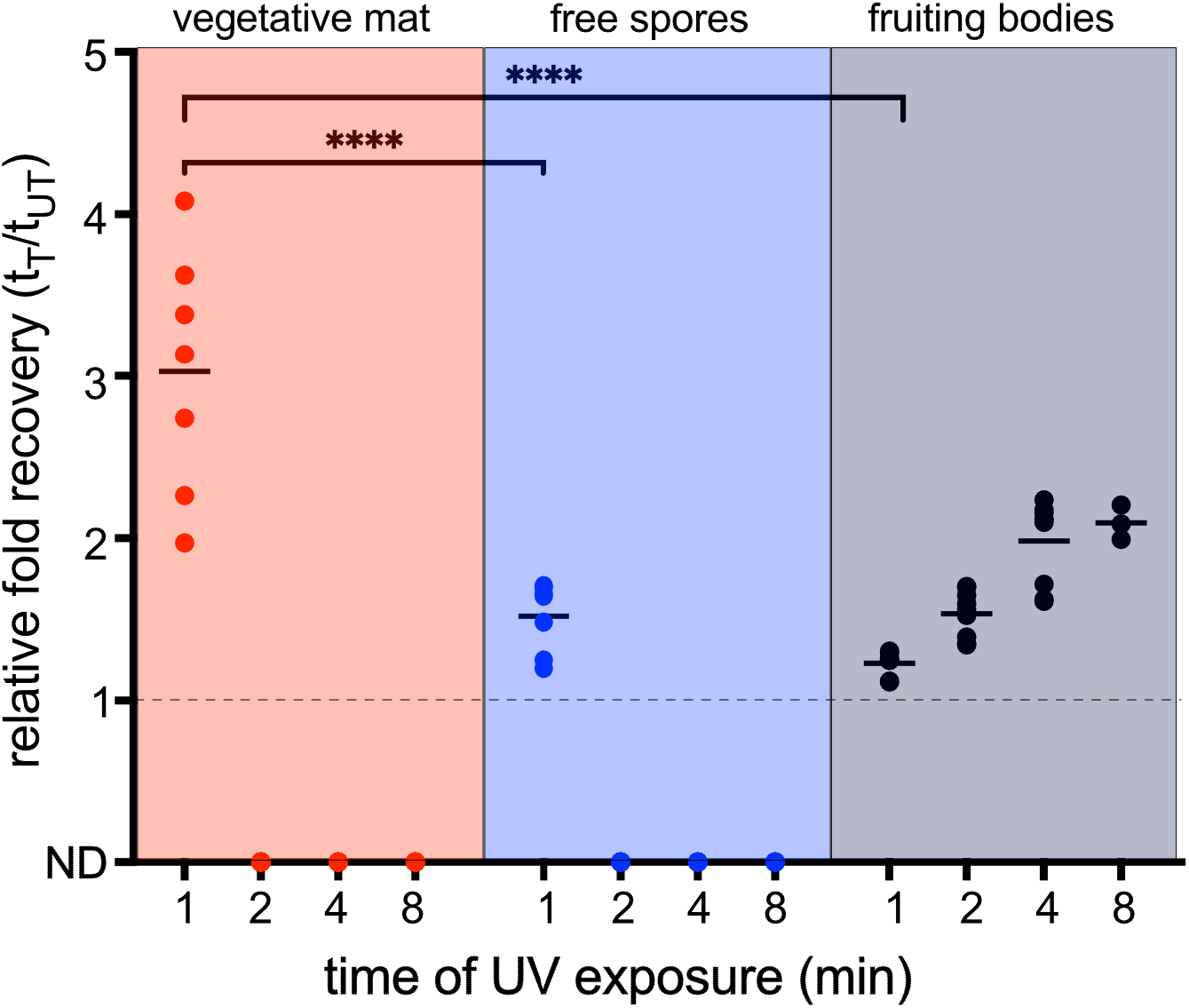
Wild-type fruiting bodies provide significant protection from UV exposure. Relative fold recovery of wild-type (DZ2) vegetative cells (red), free spores (blue), or fruiting bodies (black) after the indicated min UV exposure. Data points are the time to recovery to 0.2 A_550_ of treated samples (t_T_) normalized to the untreated sample (t_UT_). Dotted line: fold recovery equivalent to untreated samples. Data was obtained from three independent biological replicates each containing triplicate samples. Mean is indicated by solid line. Statistical significance was determined by performing two-way ANOVA followed by Tukey9s multiple comparison test (\*\*\*\**p* <0.0001). All timepoints comparisons are not significant unless indicated. t_T_ (treated time) is the time to A_550_ of 0.2 after treatment. t_UT_ (untreated time) is the time to A_550_ of 0.2 without treatment. ND: growth to 0.2 A_550_ was not detected by 72 hours.

### Sporulation is sufficient for efficent recovery from desiccation

We next set out to examine relative recovery of the cell states after exposure to increasing periods of desiccation. For this assay, vegetative mats, free spores, or fruiting bodies were desiccated for 1, 2, 4, 8, 16, or 32 days at 32 ^°^C, and recovery was monitored as described previously. Independent biological replicates demonstrated similar recovery patterns indicating that recovery time was consistent with fraction of the population that survived desiccation (data not shown).

We observed that vegetative mats failed to recover after only one day of desiccation, but free spores displayed no significant loss of recovery for up to eight days of desiccation (Fig. 4). Prolonged desiccation for 16 and 32 days only increased the recovery time relative to the untreated spores up to 2-fold (Fig. 4). Fruiting bodies also provided full protection for at least four days of desiccation. Surprisingly, however, after 16 or 32 days of desiccation, fruiting bodies took significantly longer to recover from desiccation than the free spores. Together, these results suggested that sporulation is sufficient to protect cells from desiccation and that assembly of spores into fruiting bodies does not provide additional protection; fruiting bodies may even be slightly detrimental after prolonged desiccation.

**FIG 4.**
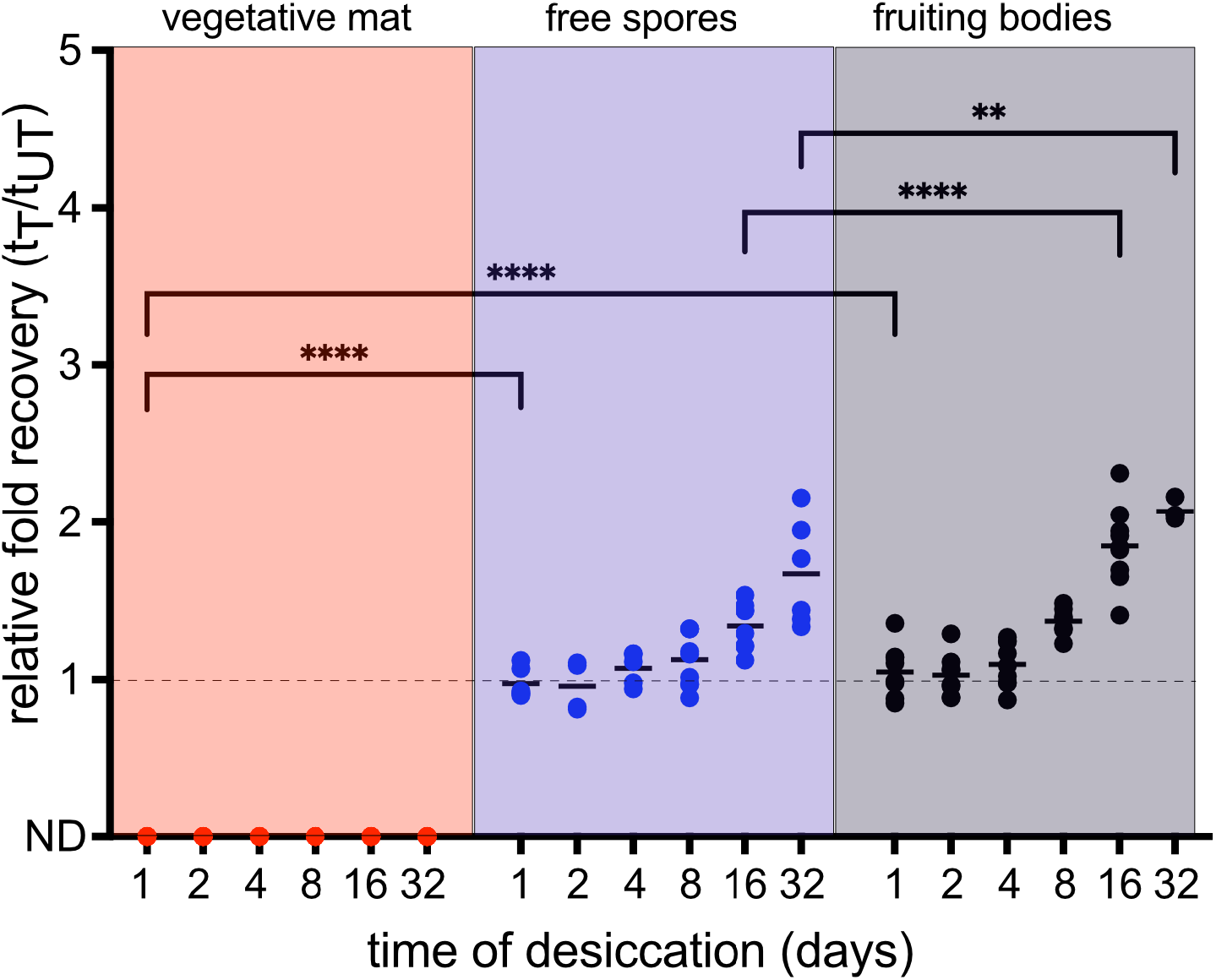
Production of spores is sufficient for protection against desiccation. Relative fold recovery of wild type (DZ2) vegetative cells (red), free spores (blue), or fruiting bodies (black) after the indicated days desiccation at 32 C. Data points are the time to recovery to 0.2 A_550_ of treated samples (t_T_) normalized to untreated samples (t_UT_). Dotted line: fold recovery equivalent to untreated samples. Data was obtained from three independent biological replicates each containing triplicate samples. Mean is indicated by solid line. Statistical significance was determined by performing two-way ANOVA followed by Tukey9s multiple comparison test (\*\*\*\**p* <0.0001). All timepoints comparisons are not significant unless indicated. t_T_ (treated time) is the time to A_550_ of 0.2 after treatment. t_UT_ (untreated time) is the time to A_550_ of 0.2 without treatment. ND: growth to 0.2 A_550_ was not detected 72 hours.

### Fruiting body morphology influences recovery from environmental insults

To examine whether fruiting body morphology is important for relative recovery from insults, we took advantage of a shallow fruiting (SF) mutant. The SF mutant lacks four genes encoding signaling systems that appear to independently inhibit accumulation of MrpC, a major transcription factor essential to induce aggregation into fruiting bodies and sporulation (32–35). Thus, in the absence of these signaling proteins, MrpC accumulates rapidly, leading to sporulation before the cells have finished moving into aggregation centers (32). As a result, the shallow and misshaped fruiting bodies were 26 ± 5 μm in height compared to 77 ± 14 μm in the wild type (Fig. 1 and 5). While the spores are viable and resistant to heat at 50 °C for 60 min and sonication (32), they appear less phase bright (Fig. 1). Regardless, these spores efficiently germinate and are a useful tool to assay whether fruiting body morphology is important for extra protection from insults.

**Fig. 5.**
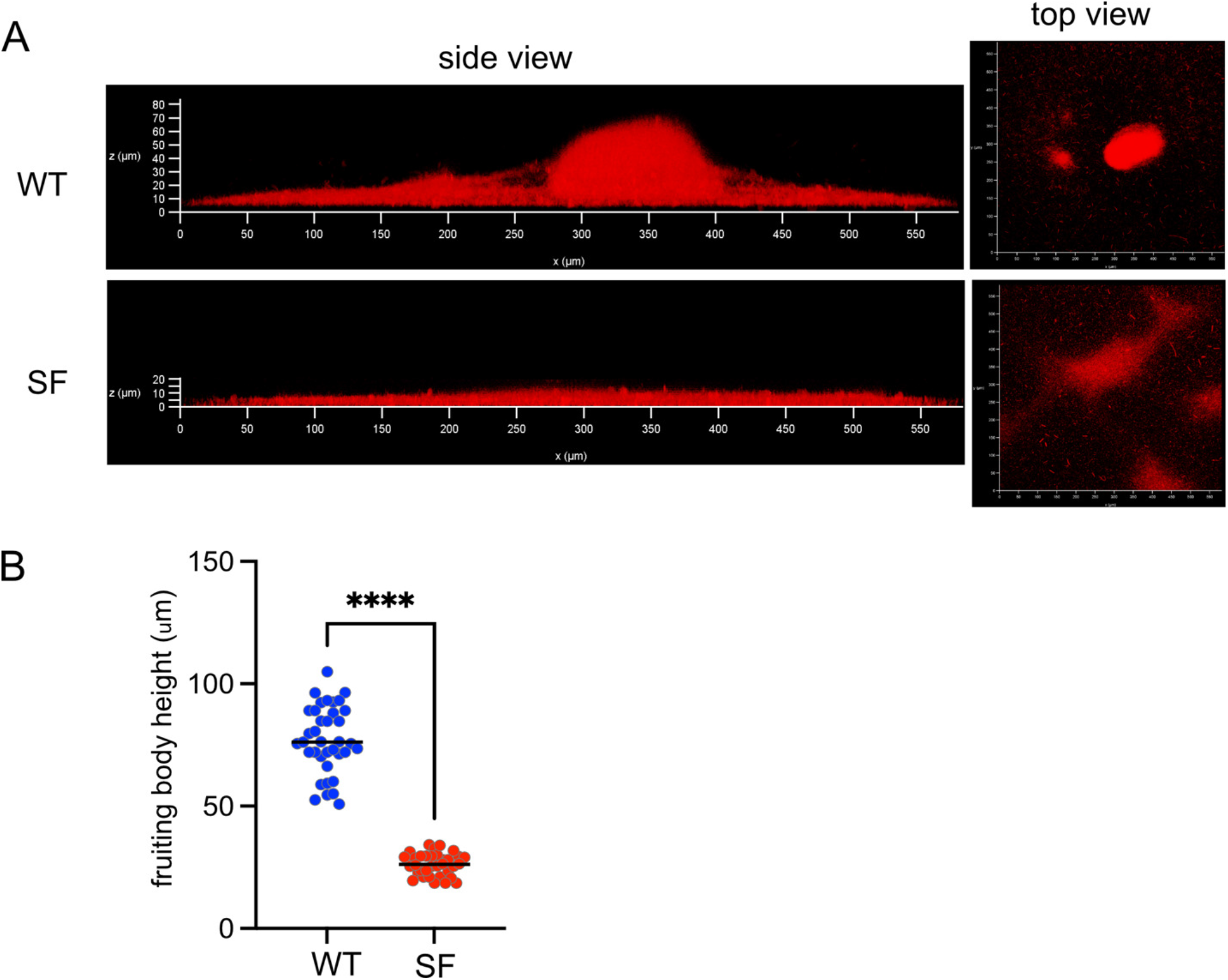
The shallow-fruiting mutant produces shallow and disorganized fruiting bodies. A) Fluorescence microscopy of wild type (DZ2) and SF (PH1054) strains expressing the fluorescent protein mCherry driven by the *pilA* promoter. Strains were developed under submerged culture for 48 hours to form fruiting bodies. B) Heights of wild type (blue) and SF mutant (red) fruiting bodies developed under submerged culture for 5 days and stained with FITC-Concanavalin A. WT, n= 37; SF, n=34 from two independent biological replicates.

Analysis of the vegetative growth pattern of the SF mutant compared to the wild type indicated that it had a similar lag time, the doubling time was reduced by one hour, the onset of stationary phase was approximately 4 hours earlier, and the death phase was not as pronounced (Table 1 and Fig. S1A and C). With respect to outgrowth of the cell states, similar lag times were observed for the vegetative mat, fruiting body, and free spore states of the SF mutant compared to the wild type, respectively.

Upon exposure to UV, we did not observe any significant differences in relative recovery times of vegetative mats or free spores between the wild type and SF mutant (Fig.6A). However, the SF mutant fruiting bodies took significantly longer (*p* <0.0001) than the wild-type fruiting bodies to recover after 1 min of UV exposure, and completely failed to recover after 2 min or more of UV exposure (Fig. 6A). These results strongly suggest that a taller, more organized fruiting body structure is required to provide significant protection against UV insults.

**FIG 6.**
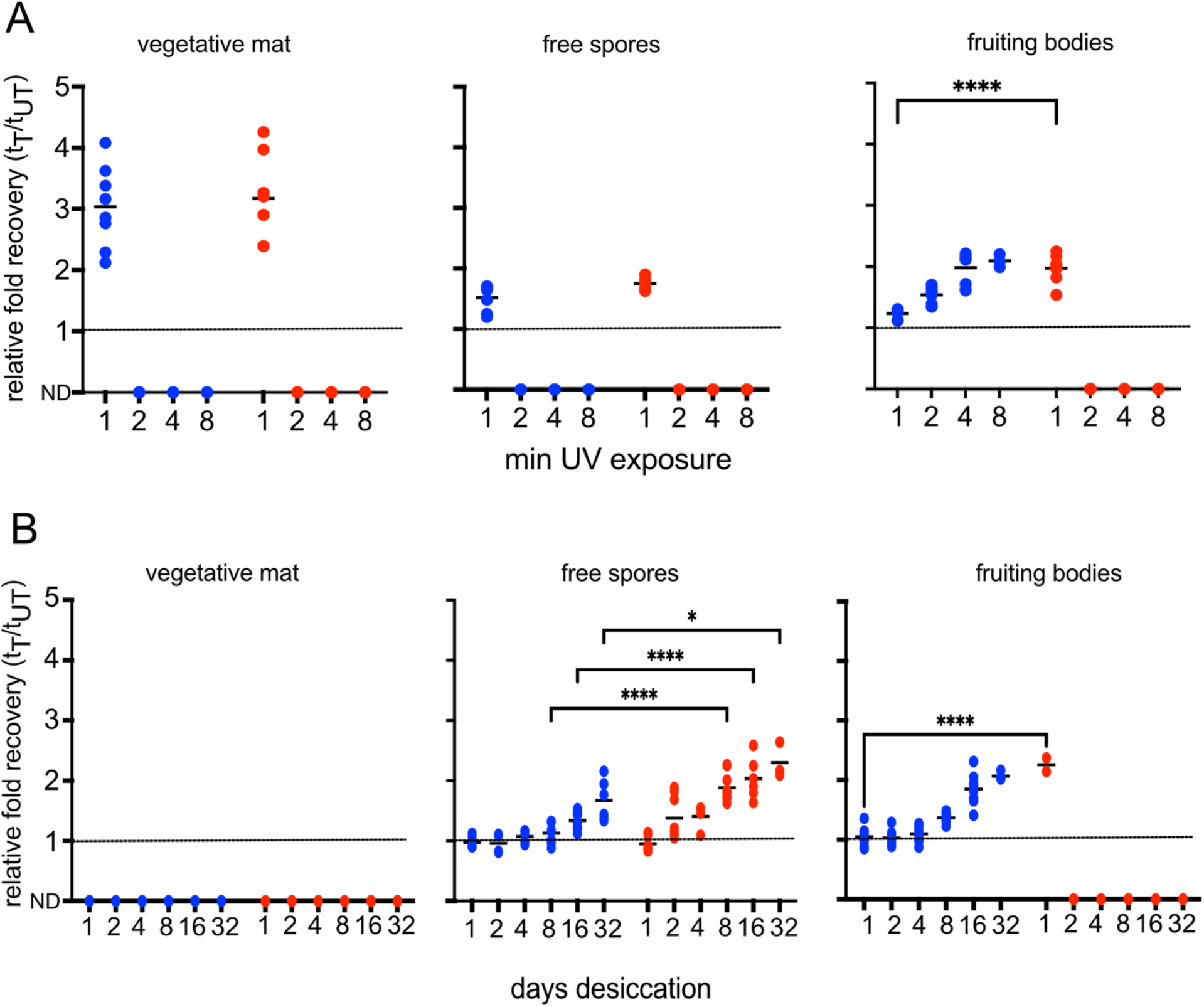
Fruiting body morphology is important for recovery from environmental insults. Relative fold recovery of wild-type (DZ2; blue) or shallow-fruiting (PH1054; red) vegetative cells, free spores, or fruiting bodies after UV exposure (A) or desiccation (B). Data points are the time to recovery to 0.2 A_550_ of treated samples (t_T_) normalized to untreated samples (t_UT_). Dotted line: fold recovery equivalent to untreated samples. Data was obtained from three independent biological replicates each containing triplicate samples. Mean is indicated by solid line. Statistical significance was determined by performing two-way ANOVA followed by Tukey9s multiple comparison test (\*\*\*\**p* <0.0001). All timepoints comparisons are not significant unless indicated. t_T_ (treated time) is the time to A_550_ of 0.2 after treatment. t_UT_ (untreated time) is the time to A_550_ of 0.2 without treatment. ND: growth to 0.2 A_550_ was not detected 72 hours.

Exposure to desiccation suggested that the SF and wild-type vegetative mats were similarly susceptible to as little as 1 day of desiccation (Fig. 6B). However, while the SF mutant and wild type free spores showed similar efficient recovery up to 4 days of desiccation, SF free spores thereafter exhibited at least 2-fold longer relative recovery times (Fig. 6B). The spores produced by the SF mutant may have some defects in desiccation resistance compared to the wild type spores. Strikingly though, in contrast to the wild type fruiting bodies, the shallow fruiting bodies did not recover after 2 days of desiccation (Fig. 6B). Interestingly, the SF mutant fruiting bodies were much more susceptible to desiccation than their free spores, which recovered even after 32 days of desiccation (Fig. 6B). Examination of the appearance of dehydrated fruiting bodies in the wild-type and SF mutant, revealed that the SF mutant fruiting bodies appeared to lose opacity within 1 day of desiccation, which correlates with failure to recover (Fig. 6). In contrast, only the edges of the wild type fruiting bodies become translucent, even after 32 days of desiccation (Fig. S5). By 2 days of desiccation, cracks occurred through the middle of the shallow fruiting mutant fruiting bodies, and around the edges of wild-type fruiting bodies (data not shown and Fig. S5), suggesting the matrix was severely dehydrated and pulling away from the surface. We suggest that dehydration of the intact ECM may hinder spore germination, with the thin SF mutant dehydrating far more quickly than the wild-type taller fruiting body.

## DISCUSSION

In this study, we examined the hypothesis that *M. xanthus* fruiting bodies protect the quiescent spores from environmental stresses. We demonstrated that spores inside wild type fruiting bodies can tolerate significantly more UV exposure than the free spores (Fig. 3). In contrast, spores themselves are highly resistant to desiccation. Fruiting bodies provide no additional protection up to ∼ 2 weeks of desiccation and actually impeded recovery after extended desiccation (Fig. 4). The haystack structure and/or height of *M. xanthus* fruiting bodies is important, because shallow disorganized fruiting bodies do not provide additional protection from UV, and significantly interfere in recovery after desiccation (Fig. 6). These results suggest that *M. xanthus* fruiting body morphology has been evolutionarily optimized to balance protection from UV and desiccation.

As part of this study, we developed a quantitative method to assess community recovery from insults rather relying on enumeration of colony forming units arising after exposure to plating single cells/spores. *M. xanthus* favors community behavior in all aspects of its lifecycle, and communities may be slightly more tolerant to insults than individual cells because of resources that may be diminished if cells are separated [*e.g.* ECM components, co-operative repair or damage dilution mechanisms (36, 37), or phenotypic heterogeneity that is destroyed upon harvesting of the community (31)].

Furthermore, recovery of the community may be enhanced by group germination (38), or by release of nutrients or eDNA from lysing cells that may promote growth and mutational repair, respectively (39). To avoid as much external manipulation of the community as possible, we therefore simply restored growth media to challenged communities and then used the time to recover to early exponential growth as a proxy for community resilience to insult. To compare the relative resistances of the cell states, we normalized the recovery times after insult to that of the respective untreated cell state.

*M. xanthus* spores isolated from fruiting bodies are well known to be more resistant to UV exposure than vegetative cells (22–24), which is at least partly due to accumulation of a small, acid-soluble protein inside the spores (24). Previous assays suggested that spores released from 10 day old fruiting bodies were up to 5.4 times more resistant to UV (15W germicidal lamp at 56 cm) than vegetative cells when assayed over the course of 90 seconds (Sudo & Dworkin, 1969). Consistently, in our assays, free spores were 2-fold more resilient to 1 min of UV exposure than vegetative mats (Fig. 3), which may be comparable given our UV dose was likely much higher. Importantly however, we demonstrate here that wild-type fruiting bodies were significantly more resistant to UV than free spores. Recovery of fruiting bodies after 1 min of UV exposure was similar to the untreated sample, and, even after 8 minutes of UV exposure, took only 2-fold longer than the untreated samples to recover. This type of UV and exposure level is biologically relevant. First, UV-B (280-320 nm) is considered the most biologically important source of UV, because it is not fully absorbed by the ozone layer and can cause direct damage to DNA via induction of pyrimidine dimers, or production of free radicals that can damage other biological processes. Second, the dose is relevant; an average June day in Florida experiences 35 KJ/m^2^ of UV-B (43). Fruiting bodies recovered from exposure to 0.08 KJ/m^2^/sec for up to 8 min, corresponding to a 32.4 KJ/m^2^ dose. We note that no recovery was observed after 16 min (64.8 KJ/m^2^) of UV exposure (data not shown), which is twice the Floridian daily dose.

Given that free spores fail to recover after more than 1 min of UV exposure (Fig. 3), we speculate that *M. xanthus* fruiting body morphology may be optimal to protect spores from environmental UV dosage. Mature wild-type fruiting bodies produced in our conditions are, on average, 77 ± 14 μm in height (Fig. 5). For reference, UV-B penetrates 10-50 μm into human epidermidis (44), and spores are likely less penetrant than layers of skin. Therefore, it is likely that UV-B does not fully penetrate to the interior of the fruiting body, and the ECM or outer layers of spores effectively shield the spores located in the interior. Consistent with this hypothesis, the shallow fruiting bodies produced by the SF mutant, with an average height that is at least 3-fold shorter than the wild type (Fig. 5), did not provide significant protection over the respective free spores (Fig. 6A). It is also possible that the fruiting body ECM, which consists of polysaccharides, proteins, eDNA, and vesicles (18–21), also contributes to protection from UV. For example, purified extracellular polysaccharides have been shown to protect from UV-B by scavenging reactive oxygen species produced during UV irradiation (45). The SF mutant does contain ECM, because the shallow fruiting bodies are efficiently labeled by the lectin Concanavalin A [which binds to some of the ECM polysaccharides (21, 46), data not shown]. However, because SF spores are produced earlier during the developmental program than in the wild type (32), the total ECM may be reduced.

Desiccation is a frequent stress experienced by environmental organisms, and extreme dehydration is lethal. Not only is water essential for metabolic activities, it also prevents aggregation of proteins, and prevents damage to membranes and DNA (47). *M. xanthus* spores exhibit two common protections against desiccation: reduced metabolic activity (48) which requires less water (49), and accumulation of trehalose (50, 51), a compatible solute that is thought to maintain protein solubility and stabilize membranes (52). In our assay, free spores released from either the wild type or SF strains could reasonably efficiently recover after desiccation for at least 32 days (Fig. 6). Surprisingly, if the spores were left in their native state, recovery was slightly (wild type) or severely (SF mutant) hindered. One difference between these two states is that free spores lack the ECM; it is disrupted during the mild sonication to release the spores from the fruiting bodies and is removed when the free spores are pelleted. We suggest severe dehydration of the fruiting body ECM leads to a “glue” that is recalcitrant to rehydration, such that germination signals failed to reach the spores and/or partially germinated cells were unable to release themselves from the glue-like matrix. This severe dehydration likely only occurs where the matrix is thin; at the edges of the wild type fruiting bodies, but throughout the shallow fruiting bodies of the SF mutant. After desiccation, these areas image as translucent and accumulate cracks associated with water loss (Fig. S5). In contrast, the centers of the wild type fruiting bodies remain opaque, and cracks do not transect the mound. Thus, while the *M. xanthus* fruiting body does not provide additional protection from desiccation spores *per se*, the haystack shape is optimized to prevent excessive dehydration of the ECM which may prevent germination.

We conclude that the *M. xanthus* fruiting bodies are not merely a means to ensure germination *en masse* into a minimal predatory population, but they play an important role in protecting the spores from environmental insults. Perhaps this is necessary because myxospores are not very hardy; they are considerably more sensitive than endospores produced by *Bacillus* and *Clostridium* species (53). This study also revealed that *M. xanthus* fruiting bodies are not equally protective against all environmental stresses, suggesting they may have evolved as an middle ground providing some resistance to several distinct stresses. However, we are interested in the possibility that *M. xanthus* can tune its fruiting body morphology to distinct environmental conditions. The SF mutant we used in this study is a deletion of numerous signal systems that regulate the timing of development (34, 54, 55). Perhaps these signaling systems monitor the environment for signals indicating rapid production of shallow fruiting body would be advantageous over haystack shaped fruiting bodies. In an extreme example, *M. xanthus* can bypass fruiting bodies entirely to produce “free” spores in response to peptidoglycan damaging agents (56), or chemical signals from other microorganisms (57).

We suspect that the variety of fruiting body morphologies produced by other myxobacteria species are optimized to provide protection from stresses encountered in different niches. *M. xanthus* is among the few myxobacteria that produce fruiting bodies of a “globular soft mucous consistency” (10); most species produce either fruiting bodies of hardened slime, or produce spores within a defined wall (sporangiole) (13, 14). It is likely that these fruiting bodies provide even more resistance to environmental insults, and it would be interesting to see if these species are enriched in more extreme environments.

## Acknowledgments

The authors gratefully acknowledge Bongsoo Lee for construction of strains PH1054 and PH2036, and Gillian Leung for construction of strain PH2037. We acknowledge past and present members of the Higgs lab for helpful discussions, and/or critical reading of the manuscript. This research was funded by a grant from the National Sciences Foundation IOS CAREER 1651921 (PIH). The funders had no role in study design, data collection and interpretation, or the decision to submit the work for publication.

## References

1. Ma B, Stirling E, Liu Y, Zhao K, Zhou J, Singh BK, Tang C, Dahlgren RA, Xu J. 2021. Soil Biogeochemical Cycle Couplings Inferred from a Function-Taxon Network. Research 2021.

2. Zakem EJ, Polz MF, Follows MJ. 2020. Redox-informed models of global biogeochemical cycles. 1. Nat Commun 11:5680.

3. Morgan AD, MacLean RC, Hillesland KL, Velicer GJ. 2010. Comparative analysis of myxococcus predation on soil bacteria. Applied and environmental microbiology, 2010/08/31 ed. 76:6920–7.

4. Nair RR, Velicer GJ. 2021. Predatory Bacteria Select for Sustained Prey Diversity. 10. Microorganisms 9:2079.

5. Mayrhofer N, Velicer GJ, Schaal KA, Vasse M. 2021. Behavioral Interactions between Bacterivorous Nematodes and Predatory Bacteria in a Synthetic Community. 7. Microorganisms 9:1362.

6. Dai W, Wang N, Wang W, Ye X, Cui Z, Wang J, Yao D, Dong Y, Wang H. 2021. Community Profile and Drivers of Predatory Myxobacteria under Different Compost Manures. Microorganisms 9:2193.

7. Petters S, Groß V, Söllinger A, Pichler M, Reinhard A, Bengtsson MM, Urich T. 2021. The soil microbial food web revisited: Predatory myxobacteria as keystone taxa? ISME J 15:2665– 2675.

8. Wang J, Wang J, Wu S, Zhang Z, Li Y. 2021. Global Geographic Diversity and Distribution of the Myxobacteria. Microbiology Spectrum 9:e00012–21.

9. Liu Y, Yao Q, Zhu H. 2019. Meta-16S rRNA Gene Phylogenetic Reconstruction Reveals the Astonishing Diversity of Cosmopolitan Myxobacteria. 11. Microorganisms 7:551.

10. Dawid W. 2000. Biology and global distribution of myxobacteria in soils. FEMS Microbiology Reviews 24:403–427.

11. Tiedje JM, Bruns MA, Casadevall A, Criddle CS, Eloe-Fadrosh E, Karl DM, Nguyen NK, Zhou J. 2022. Microbes and Climate Change: a Research Prospectus for the Future. mBio 13:e00800–22.

12. Cavicchioli R, Ripple WJ, Timmis KN, Azam F, Bakken LR, Baylis M, Behrenfeld MJ, Boetius A, Boyd PW, Classen AT, Crowther TW, Danovaro R, Foreman CM, Huisman J, Hutchins DA, Jansson JK, Karl DM, Koskella B, Mark Welch DB, Martiny JBH, Moran MA, Orphan VJ, Reay DS, Remais JV, Rich VI, Singh BK, Stein LY, Stewart FJ, Sullivan MB, van Oppen MJH, Weaver SC, Webb EA, Webster NS. 2019. Scientists’ warning to humanity: microorganisms and climate change. 9. Nat Rev Microbiol 17:569–586.

13. Reichenbach H. 1993. Biology of the Myxobacteria: Ecology and Taxonomy, p. 13–62. In Dworkin, Martin, Kaiser, Dale (ed.), Myxobacteria II. American Society for Microbiology.

14. Reichenbach H. 1984. Myxobacteria: A Most Peculiar Group of Social Prokaryotes, p. 1–50. In Rosenberg, E (ed.), Myxobacteria, development and cell interactions. Springer-Verlag.

15. Thiery S, Kaimer C. 2020. The Predation Strategy of Myxococcus xanthus. Front Microbiol, 2020/02/06 ed. 11:2.

16. Phillips KE, Akbar S, Stevens DC. 2022. Concepts and conjectures concerning predatory performance of myxobacteria. Frontiers in Microbiology 13.

17. Rosenberg E, Keller KH, Dworkin M. 1977. Cell density-dependent growth of Myxococcus xanthus on casein. Journal of bacteriology, 1977/02/01 ed. 129:770–7.

18. Behmlander RM, Dworkin M. 1994. Biochemical and structural analyses of the extracellular matrix fibrils of Myxococcus xanthus. Journal of bacteriology, 1994/10/01 ed. 176:6295–303.

19. Pathak DT, Wei X, Bucuvalas A, Haft DH, Gerloff DL, Wall D. 2012. Cell contact-dependent outer membrane exchange in myxobacteria: genetic determinants and mechanism. PLoS Genet, 2012/04/19 ed. 8:e1002626.

20. Hu W, Li L, Sharma S, Wang J, McHardy I, Lux R, Yang Z, He X, Gimzewski JK, Li Y, Shi W. 2012. DNA builds and strengthens the extracellular matrix in Myxococcus xanthus biofilms by interacting with exopolysaccharides. PloS one, 2013/01/10 ed. 7:e51905.

21. Islam ST, Vergara Alvarez I, Saidi F, Guiseppi A, Vinogradov E, Sharma G, Espinosa L, Morrone C, Brasseur G, Guillemot JF, Benarouche A, Bridot JL, Ravicoularamin G, Cagna A, Gauthier C, Singer M, Fierobe HP, Mignot T, Mauriello EMF. 2020. Modulation of bacterial multicellularity via spatio-specific polysaccharide secretion. PLoS Biol, 2020/06/10 ed. 18:e3000728.

22. Sudo SZ, Dworkin M. 1969. Resistance of Vegetative Cells and Microcysts of Myxococcus xanthus. J Bacteriol 98:883–887.

23. Tengra FK, Dahl JL, Dutton D, Caberoy NB, Coyne L, Garza AG. 2006. CbgA, a protein involved in cortex formation and stress resistance in Myxococcus xanthus spores. Journal of bacteriology, 2006/09/26 ed. 188:8299–302.

24. Dahl JL, Fordice D. 2011. Small acid-soluble proteins with intrinsic disorder are required for UV resistance in Myxococcus xanthus spores. Journal of bacteriology, 2011/04/26 ed. 193:3042–8.

25. Galperin MY. 2013. Genome Diversity of Spore-Forming Firmicutes. Microbiol Spectr 1:TBS-0015–2012.

26. Campos JM, Zusman DR. 1975. Regulation of development in Myxococcus xanthus: effect of 3’:5’-cyclic AMP, ADP, and nutrition. Proc Natl Acad Sci USA 72:518–522.

27. Muller FD, Treuner-Lange A, Heider J, Huntley SM, Higgs PI. 2010. Global transcriptome analysis of spore formation in Myxococcus xanthus reveals a locus necessary for cell differentiation. BMC genomics, 2010/04/28 ed. 11:264.

28. Wood JL, Osman A, Wade SA. 2019. An efficient, cost-effective method for determining the growth rate of sulfate-reducing bacteria using spectrophotometry. MethodsX 6:2248– 2257.

29. Lee B, Schramm A, Jagadeesan S, Higgs PI. 2010. Two-component systems and regulation of developmental progression in Myxococcus xanthus. Methods in enzymology, 2010/10/16 ed. 471:253–78.

30. Higgs PI, Merlie Jr JP. 2008. Myxococcus xanthus: Cultivation, Motility, and Development, p. 465–478. In David Whitworth (ed.), Myxobacteria: Multicellularity and Differentiation. ASM Press.

31. Lee B, Holkenbrink C, Treuner-Lange A, Higgs PI. 2012. Myxococcus xanthus developmental cell fate production: heterogeneous accumulation of developmental regulatory proteins and reexamination of the role of MazF in developmental lysis. Journal of bacteriology, 2012/04/12 ed. 194:3058–68.

32. 32. Lee B, Higgs PI. 2009. The role of negative regulators in coordination fo the Myxococcus xanthus developmental program. PhD Thesis University of Marburg.

33. Higgs PI, Jagadeesan S, Mann P, Zusman DR. 2008. EspA, an orphan hybrid histidine protein kinase, regulates the timing of expression of key developmental proteins of Myxococcus xanthus. Journal of bacteriology, 2008/04/09 ed. 190:4416–26.

34. Schramm A, Lee B, Higgs PI. 2012. Intra- and interprotein phosphorylation between two-hybrid histidine kinases controls Myxococcus xanthus developmental progression. The Journal of biological chemistry, 2012/06/05 ed. 287:25060–72.

35. Sun H, Shi W. 2001. Genetic studies of mrp, a locus essential for cellular aggregation and sporulation of Myxococcus xanthus. Journal of bacteriology, 2001/07/24 ed. 183:4786–95.

36. Dey A, Wall D. 2014. A genetic screen in Myxococcus xanthus identifies mutants that uncouple outer membrane exchange from a downstream cellular response. Journal of bacteriology, 2014/10/01 ed. 196:4324–32.

37. Vassallo CN, Wall D. 2016. Tissue repair in myxobacteria: A cooperative strategy to heal cellular damage. Bioessays 38:306–315.

38. Pande S, Pérez Escriva P, Yu Y-TN, Sauer U, Velicer GJ. 2020. Cooperation and Cheating among Germinating Spores. Current Biology 30:4745–4752.e4.

39. Wang J, Hu W, Lux R, He X, Li Y, Shi W. 2011. Natural transformation of Myxococcus xanthus. Journal of bacteriology, 2011/03/08 ed. 193:2122–32.

40. Kottel RH, Bacon K, Clutter D, White D. 1975. Coats from Myxococcus xanthus: characterization and synthesis during myxospore differentiation. Journal of Bacteriology 124:550–557.

41. Inouye M, Inouye S, Zusman DR. 1979. Biosynthesis and self-assembly of protein S, a development-specific protein of Myxococcus xanthus. Proc Natl Acad Sci U S A 76:209–213.

42. Abhyankar W, Pandey R, Ter Beek A, Brul S, de Koning LJ, de Koster CG. 2015. Reinforcement of Bacillus subtilis spores by cross-linking of outer coat proteins during maturation. Food Microbiol 45:54–62.

43. Lee-Taylor J, Madronich S, Fischer C, Mayer B. 2010. A Climatology of UV Radiation, 1979– 2000, 65S–65N, p. 1–20. In Gao, W, Slusser, JR, Schmoldt, DL (eds.), UV Radiation in Global Climate Change: Measurements, Modeling and Effects on Ecosystems. Springer, Berlin, Heidelberg.

44. Meinhardt M, Krebs R, Anders A, Heinrich U, Tronnier H. 2008. Wavelength-dependent penetration depths of ultraviolet radiation in human skin. JBO 13:044030.

45. Peng S, Guo C, Wu S, Duan Z. 2022. Isolation, characterization and anti-UVB irradiation activity of an extracellular polysaccharide produced by Lacticaseibacillus rhamnosus VHPriobi O17. Heliyon 8:e11125.

46. Ducret A, Fleuchot B, Bergam P, Mignot T. 2013. Direct live imaging of cell-cell protein transfer by transient outer membrane fusion in Myxococcus xanthus. eLife, 2013/07/31 ed. 2:e00868.

47. Aung K, Jiang Y, He SY. 2018. The role of water in plant–microbe interactions. The Plant Journal 93:771–780.

48. Watson BF, Dworkin M. 1968. Comparative Intermediary Metabolism of Vegetative Cells and Microcysts of Myxococcus xanthus. Journal of Bacteriology 96:1465–1473.

49. Rittershaus ESC, Baek S, Sassetti CM. 2013. The Normalcy of Dormancy. Cell Host Microbe 13:643–651.

50. McBride MJ, Zusman DR. 1989. Trehalose accumulation in vegetative cells and spores of Myxococcus xanthus. Journal of bacteriology, 1989/11/01 ed. 171:6383–6.

51. Kimura Y, Kawasaki S, Tuchimoto R, Tanaka N. 2014. Trehalose biosynthesis in Myxococcus xanthus under osmotic stress and during spore formation. Journal of biochemistry, 2013/10/08 ed. 155:17–24.

52. Potts M. 1994. Desiccation tolerance of prokaryotes. Microbiological Reviews 58:755–805.

53. Nicholson WL, Fajardo-Cavazos P, Rebeil R, Slieman TA, Riesenman PJ, Law JF, Xue Y. 2002. Bacterial endospores and their significance in stress resistance. Antonie Van Leeuwenhoek 81:27–32.

54. Jagadeesan S, Mann P, Schink CW, Higgs PI. 2009. A novel “four-component” two-component signal transduction mechanism regulates developmental progression in Myxococcus xanthus. The Journal of biological chemistry, 2009/06/19 ed. 284:21435–45.

55. Rasmussen AA, Sogaard-Andersen L. 2003. TodK, a putative histidine protein kinase, regulates timing of fruiting body morphogenesis in Myxococcus xanthus. Journal of bacteriology, 2003/09/02 ed. 185:5452–64.

56. O’Connor KA, Zusman DR. 1997. Starvation-independent sporulation in Myxococcus xanthus involves the pathway for beta-lactamase induction and provides a mechanism for competitive cell survival. Molecular microbiology, 1997/05/01 ed. 24:839–50.

57. Marcos-Torres FJ, Volz C, Muller R. 2020. An ambruticin-sensing complex modulates Myxococcus xanthus development and mediates myxobacterial interspecies communication. Nat Commun, 2020/11/06 ed. 11:5563.

